# Comprehensive Atlas of Cancer-Type-Specific Molecular Features from Comparative Analysis of TCGA Data

**DOI:** 10.1101/2025.10.26.684620

**Authors:** Yeu-Guang Tung

## Abstract

Cancer-type-specific molecular alterations often reflect the unique biological context of their tissue of origin and are more likely to represent relevant drivers rather than passenger events. We analyzed data from The Cancer Genome Atlas (TCGA) spanning 39 cancer types and 6 molecular platforms to create a comprehensive atlas of cancer-type-specific molecular features through unified comparative analyses. Simple nucleotide variation analysis characterized cancer-type-specific gene mutations and revealed heterogeneity among mutated genes within shared pathways. Copy number variation analysis characterized cancer-type-specific amplifications and deletions and demonstrated synergistic interactions between gene deletions and mutations. DNA methylation analysis identified candidate hypermethylated genes alongside well-established targets. Transcriptome profiling analysis revealed cancer-type-specific pathway enrichment reflecting tissue-of-origin functions or novel associations. Multiomics clustering analysis identified multi-cancer clusters and revealed consistent patterns across molecular platforms. These findings provide insights into cancer-type-specific molecular features and offer comprehensive visualizations as a reference resource for clinical application and hypothesis generation.

## Introduction

Cancer is a leading cause of morbidity and mortality worldwide, with an estimated 20 million new cases and 9.7 million deaths occurring globally in 2022 [1]. To better understand cancer biology and improve diagnosis and treatment, advances in molecular biology have revealed diverse molecular characteristics of cancer beyond traditional gross, microscopic, and immunohistochemical features [2]. The Cancer Genome Atlas (TCGA), a landmark project spanning more than 10 years and analyzing more than 11,000 cases across 33 tumor types, has provided comprehensive molecular data of various cancers across multiple platforms [3]. Individual TCGA studies characterized specific genomic, epigenomic, transcriptomic, and proteomic landscapes of different cancers [4] [5] [6], while TCGA pan-cancer studies investigated shared cell-of-origin patterns, oncogenic processes, and signaling pathways across cancers [7] [8] [9].

Despite the tremendous contributions of prior TCGA studies, there remains a lack of research focused on cancer-type-specific molecular features identified through comparative analysis across the entire dataset. Specifically, individual TCGA studies investigated cancer-type-specific features by analyzing single cancer types, each employing distinct analytical pipelines and statistical cutoffs to identify significant alterations, making direct comparison across cancer types problematic. Conversely, TCGA pan-cancer studies primarily investigated common features by analyzing multiple cancer types, aiming to obtain generalized insights applicable to the majority of cancers rather than characterizing molecular differences between distinct cancer types. Consequently, few studies have investigated cancer-type-specific features by analyzing multiple cancer types—that is, using unified comparative approaches across the entire dataset to distinguish what is unique to each cancer type. Molecular features that are significantly associated with specific cancer types are crucial, as they may indicate that these molecular alterations are biologically relevant drivers rather than randomly occurring passengers. They may also reflect that the organs and cells of the primary site have unique biological contexts that confer susceptibility to specific molecular alterations.

In this work, we present a comprehensive atlas of cancer-type-specific molecular features from comparative analysis of TCGA data. We employed minimally processed data publicly available from TCGA rather than relying on pre-processed summary datasets. We implemented a uniform analysis pipeline with consistent statistical cutoffs, enabling robust inter-cancer-type comparisons. We analyzed all available cancer types across all available molecular platforms, ensuring a complete pan-cancer multi-omics viewpoint. Through this systematic approach, we gained novel insights into cancer-type-specific molecular features, and provided clear visualizations as a resource for reference and hypothesis generation.

## Results

### Cohort and Dataset

In this pan-cancer multi-omics study, we analyzed data from The Cancer Genome Atlas (TCGA) [3] to characterize cancer type-specific molecular alterations. We established cohorts for 39 distinct malignancies representing major anatomical sites and histological subtypes based on histopathological diagnosis. We incorporated all available TCGA data from 6 different molecular platforms. In total, 11,073 individuals were analyzed.

The 39 cancer types included: Lung squamous cell carcinoma (LUSC, n=503), lung adenocarcinoma (LUAD, n=488), esophageal squamous cell carcinoma (ESCC, n=96), esophageal adenocarcinoma (EAC, n=89), gastric adenocarcinoma (GAC, n=443), colorectal adenocarcinoma (CRC, n=627), hepatocellular carcinoma (HCC, n=369), cholangiocarcinoma (CCA, n=48), pancreatic ductal adenocarcinoma (PDAC, n=176), head and neck squamous cell carcinoma (HNSCC, n=528), thyroid papillary carcinoma (PTC, n=504), adrenocortical carcinoma (ACC, n=92), pheochromocytoma and paraganglioma (PPGL, n=179), clear cell renal cell carcinoma (ccRCC, n=523), papillary renal cell carcinoma (pRCC, n=291), chromophobe renal cell carcinoma (chRCC, n=113), bladder urothelial carcinoma (BUC, n=410), prostate adenocarcinoma (PCa, n=500), testicular germ cell tumor (TGCT, n=247), cervical squamous cell carcinoma (CSCC, n=254), cervical adenocarcinoma (CAC, n=52), endometrial carcinoma (EC, n=546), uterine carcinosarcoma (UCS, n=56), ovarian serous carcinoma (OSC, n=587), breast invasive ductal carcinoma (IDC, n=816), breast invasive lobular carcinoma (ILC, n=209), cutaneous melanoma (SKCM, n=470), uveal melanoma (UM, n=79), liposarcoma (LPS, n=62), leiomyosarcoma (LMS, n=104), undifferentiated pleomorphic sarcoma (UPS, n=49), pleural mesothelioma (MPM, n=87), astrocytoma (AC, n=195), oligoastrocytoma (OAC, n=131), oligodendroglioma (ODG, n=190), glioblastoma (GBM, n=599), acute myeloid leukemia (AML, n=200), diffuse large B-cell lymphoma (DLBCL, n=48), and thymoma (THYM, n=113).

The 6 molecular platforms included: Simple nucleotide variation (SNV, n=9,922), copy number variation (CNV, n=10,816), DNA methylation (Meth, n=10,674), mRNA expression (mRNA, n=10,260), miRNA expression (miRNA, n=10,311), and protein expression (Prot, n=7,650). For DNA methylation analysis, we included negative controls from normal adjacent tissues in TCGA (n=1,075) and normal leukocytes from the GSE35069 dataset (n=60) [10].

To provide context for our comparative analysis, we briefly summarized the characteristic molecular alterations of various cancer types identified in previous studies. Lung squamous cell carcinoma is characterized by *TP53* mutation, *CDKN2A* inactivation, *NFE2L2*/*KEAP1* mutation, and *SOX2*/*TP63* amplification [11]. Lung adenocarcinoma is characterized by *TP53* mutation, *CDKN2A* inactivation, *KRAS* mutation, *EGFR* mutation, and *ROS1*/*ALK* fusion [12]. Esophageal cancer is characterized by *TP53* mutation, *CDKN2A* inactivation, *CCND1* amplification and *TP63*/*SOX2* amplification (squamous cell carcinoma), and *ERBB2* amplification (adenocarcinoma) [13]. Gastric adenocarcinoma is characterized by *TP53* mutation (CIN), *MLH1* silencing (MSI), *CDH1* mutation (GS), and *PIK3CA* mutation (EBV) [14]. Colorectal adenocarcinoma is characterized by *APC* mutation, *TP53* mutation, and *KRAS* mutation (CIN), and *MLH1* silencing (MSI) [5]. Hepatocellular carcinoma is characterized by *TP53* mutation, *CTNNB1* mutation, *TERT* promoter mutation, and *CDKN2A* deletion [15]. Cholangiocarcinoma is characterized by *IDH1/2* mutation, *FGFR2* fusion, *BAP1* mutation, and *PBRM1* mutation [16]. Pancreatic ductal adenocarcinoma is characterized by *KRAS* mutation, *TP53* mutation, *CDKN2A* inactivation, and *SMAD4* inactivation [17]. Head and neck squamous cell carcinoma is characterized by *TP53* mutation and *CDKN2A* inactivation (HPV-negative), and *PIK3CA* mutation (HPV-positive) [18]. Papillary thyroid carcinoma is characterized by *BRAF* mutation, *N/H/KRAS* mutation, and *RET* fusion [19]. Adrenocortical carcinoma is characterized by *TP53* mutation, *CTNNB1* mutation, and *ZNRF3* deletion [20]. Pheochromocytoma/Paraganglioma is characterized by *NF1* mutation, *HRAS* mutation, *RET* mutation, *SDHB/D* mutation, and *VHL* mutation [21]. Clear cell renal cell carcinoma is characterized by *VHL* mutation, *PBRM1* mutation, *SETD2* mutation, and *BAP1* mutation [22]. Papillary renal cell carcinoma is characterized by *MET* mutation (type 1) and *CDKN2A* inactivation (type 2) [23]. Chromophobe renal cell carcinoma is characterized by *TP53* mutation and *PTEN* mutation [24]. Urothelial carcinoma of bladder is characterized by *TP53* mutation, *PIK3CA* mutation, *FGFR3* mutation, *CDKN2A* deletion, and *RB1* inactivation [25] [26]. Prostate adenocarcinoma is characterized by *TMPRSS2*-*ERG*/*ETV1*/*ETV4* fusion, *SPOP* mutation, *PTEN* inactivation, and *TP53* mutation [27]. Testicular germ cell tumor is characterized by *KIT* mutation and *KRAS* activation (seminoma), and *KRAS* amplification (non-seminoma) [28]. Cervical cancer is characterized by *PIK3CA* activation (squamous cell carcinoma), *ERBB2* activation and *KRAS* mutation (adenocarcinoma) [29]. Endometrial carcinoma is characterized by *PTEN* mutation (endometrioid, CN-low), *MLH1* silencing (endometrioid, MSI), *POLE* mutation (endometrioid, POLE), and *TP53* mutation (serous, CN-high) [30]. Uterine carcinosarcoma is characterized by *TP53* mutation, *FBXW7* mutation, and *PIK3CA* mutation [31]. Ovarian serous carcinoma is characterized by *TP53* mutation and *BRCA1/2* mutation [32]. Breast cancer is characterized by *PIK3CA* mutation (luminal A/B), *GATA3* mutation (luminal A, IDC), *CDH1* mutation (luminal A, ILC), *ERBB2* amplification (HER2-enriched), and *TP53* mutation (basal-like) [4] [33]. Cutaneous melanoma is characterized by *BRAF* mutation, *NRAS* mutation, and *NF1* mutation [34]. Uveal melanoma is characterized by *GNAQ*/*GNA11* mutation, *BAP1* mutation (monosomy 3), and *SF3B1* mutation (disomy 3) [35]. Soft tissue sarcoma is characterized by *TP53* mutation, *ATRX* mutation, *RB1* inactivation, *CDKN2A* deletion, *MDM2* and *CDK4* amplification (dedifferentiated liposarcoma), *MYOCD* amplification (leiomyosarcoma), and *VGLL3* amplification (undifferentiated pleomorphic sarcoma and myxofibrosarcoma) [36]. Pleural mesothelioma is characterized by *BAP1* inactivation, *NF2* inactivation, and *CDKN2A* deletion [37]. Lower-grade glioma is characterized by *IDH1/2* mutation, 1p/19q codeletion, *CIC* and *FUBP1* mutation (*IDH*-mut, 1p/19q-del), *TP53* and *ATRX* mutation (*IDH*-mut, no 1p/19q-del), and *CDKN2A* deletion and *EGFR* activation (*IDH*-wild) [38]. Glioblastoma is characterized by *CDKN2A* deletion, *EGFR* activation, *PTEN* inactivation, and *TP53* mutation [6] [39]. Acute myeloid leukemia is characterized by *NPM1* mutation, *FLT3* mutation, *DNMT3A* mutation, *IDH1/2* mutation, *PML*-*RARA* fusion, *CBFB*-*MYH11* fusion, and *RUNX1*-*RUNX1T1* fusion [40]. Diffuse large B-cell lymphoma is characterized by *MYD88* mutation and *CD79B* mutation (ABC subtype), *BCL2* fusion and *EZH2* mutation (GCB subtype), and *BCL6* fusion and *TP53* mutation (mixed) [41]. Thymoma is characterized by *GTF2I* mutation [42].

### Simple Nucleotide Variation Analysis

We analyzed the simple nucleotide variation data to characterize cancer-type-specific gene mutations and mutational hotspots. Mutations in tumor-related genes were summarized for each cancer type, where the tumor-related genes analyzed in this study included 460 genes curated by the Japanese Cancer Genome Atlas (JCGA) [43] with the manual addition of *FOXP1, PAX8, LATS1, LATS2, GTF2I*, and *MYOCD*. The waterfall plots display the mutation distribution of the most frequently mutated tumor-related genes in each cancer type (Figure 1). The bubble plots show gene mutation frequencies (left) and hotspot frequencies (right) for genes with high mutation frequencies in any cancer type and their prevalent mutational hotspots (Figure 2).

**Figure 1:**
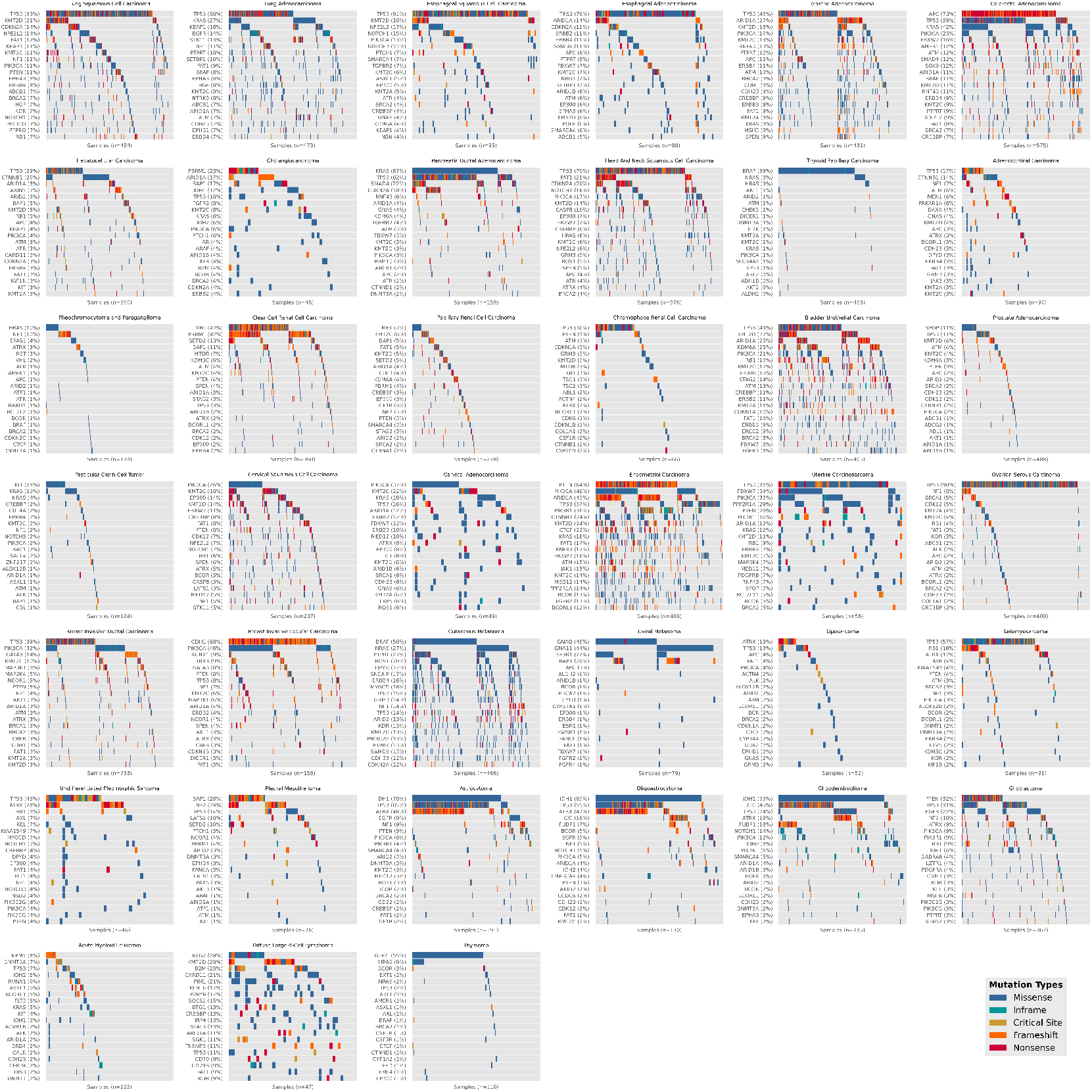
Mutational landscapes across cancer types. Mutation distribution of tumor-related genes across cancer types. Each waterfall plot displays the mutation status of samples within a cancer type. Color coding represents mutation types: missense mutations (blue), in-frame indels (teal), critical site mutations (yellow), frameshift indels (orange), and nonsense mutations (red). Genes are ordered by mutation frequency. Numbers following gene names indicate mutation frequencies; numbers below each waterfall plot indicate sample sizes.

**Figure 2:**
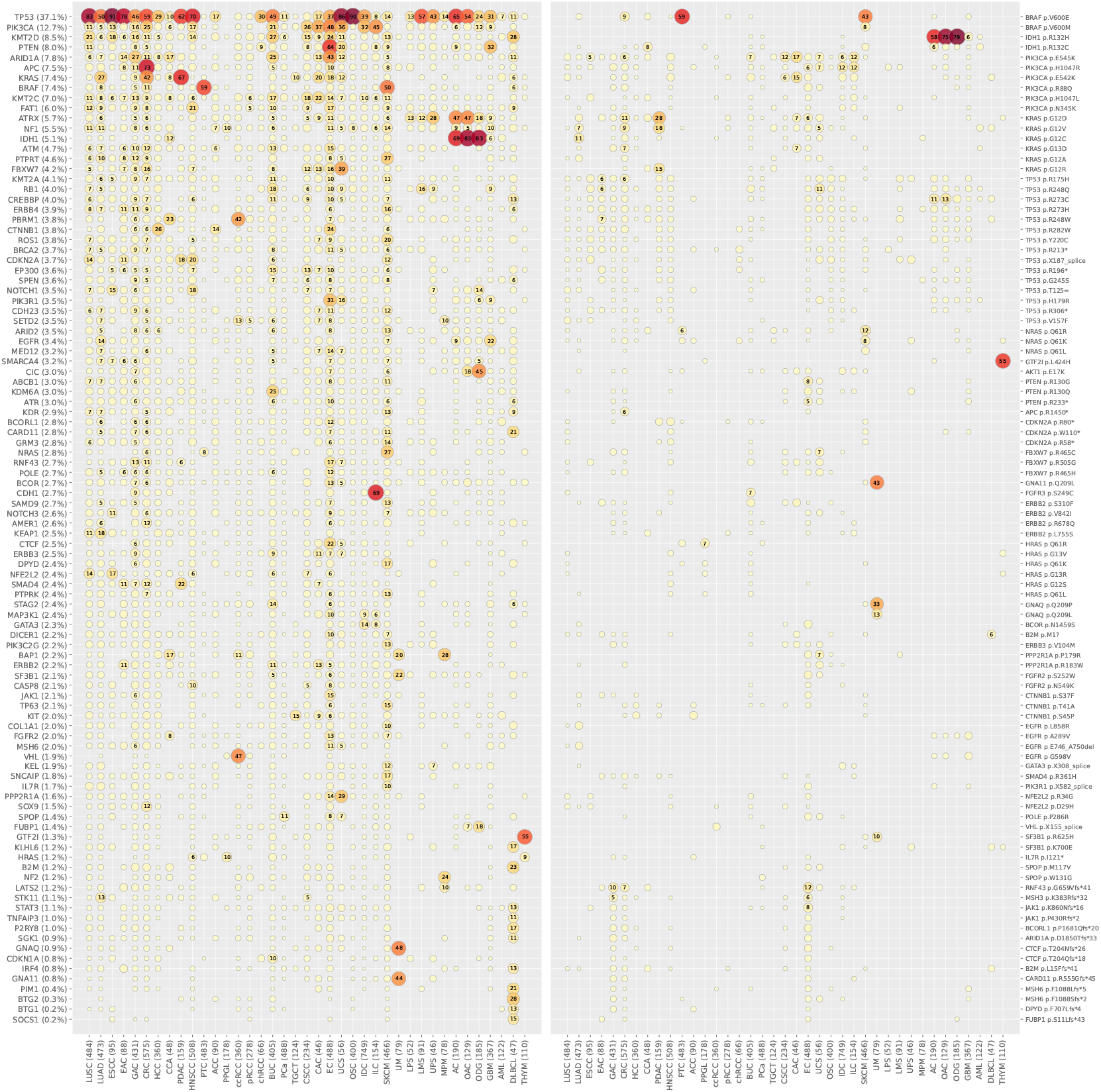
Gene mutations and hotspots across cancer types. Gene mutation frequencies (left) and hotspot frequencies (right) for tumor-related genes across cancer types. Bubble size and color intensity represent alteration frequency. Genes and hotspots are ordered by overall mutation frequency, with hotspots from the same gene grouped together, and frameshift mutations positioned at the end of the plot. Numbers following gene names indicate overall mutation frequencies; numbers following cancer type names indicate sample sizes.

The majority of cancer types exhibited widespread mutations across various genes, with *TP53* typically being the most frequently mutated gene. Notable exceptions include CRC (*APC*), PDAC (*KRAS*), ccRCC (*VHL*), CSCC and CAC (*PIK3CA*), EC (*PTEN*), ILC (*CDH1*), SKCM (*BRAF*), and AC, OAC, and ODG (*IDH1*). Several cancer types demonstrated highly selective mutations in specific genes, including PTC (*BRAF*), UM (*GNAQ*/*GNA11*), and THYM (*GTF2I*). Cancer types with a large proportion of samples lacking mutations typically rely on oncogenic processes other than somatic mutations, including PCa (ETS family fusion), TGCT (chromosome 12p gain), LPS (*MDM2*/*CDK4* amplification), and PPGL (germline mutation). Interestingly, the nature of tumor-related genes can also be inferred from the mutational landscape. Oncogenes, including *PIK3CA, KRAS*, and *BRAF*, predominantly exhibited missense mutations. Tumor suppressor genes, including *TP53, PTEN*, and *APC*, demonstrated a mixture of missense, frameshift, and nonsense mutations. Genes susceptible to microsatellite instability (MSI), including *RNF43*, showed frequent frameshift mutations, while *EGFR* showed unusually frequent in-frame indels in LUAD.

*TP53* was the most frequently altered gene, mutated in 37% of all tumors. While 29 out of 39 cancer types showed *TP53* mutation in more than 10% of cases, there was a notable pattern favoring squamous cell carcinomas over adenocarcinomas and other cancer types, with ESCC, LUSC, and HNSCC showing higher mutation frequencies than LUAD, GAC, CRC, and PDAC. Additionally, OSC, UCS, and EAC showed exceptionally high mutation frequencies. *PIK3CA* was the most frequently altered oncogene, with mutational hotspots at E545K, H1047R, and E542K. Notably, the cancer types with the highest *PIK3CA* mutation frequencies were predominantly gynecologic malignancies, including EC, UCS, CSCC, CAC, IDC, and ILC. *PTEN* exhibited a selective alteration pattern, with significant mutations in EC, followed by GBM and UCS. *PIK3R1* also showed a selective alteration pattern in EC, followed by UCS. Despite sharing the PI3K/AKT pathway, *PTEN* and *PIK3R1* demonstrated much more restricted mutation distributions primarily confined to EC, compared to *PIK3CA*’s widespread occurrence across multiple gynecologic cancers. *KRAS* was a frequently altered oncogene with mutational hotspots at G12D, G12V, and G12C, while PDAC additionally exhibited a specific G12R hotspot. *KRAS* mutations showed selective enrichment in adenocarcinomas, particularly PDAC, CRC, LUAD, and CAC. Notably, upper gastrointestinal adenocarcinomas (EAC and GAC) showed significantly lower *KRAS* mutation frequencies. *BRAF* exhibited highly selective mutations in PTC and SKCM, predominantly at mutational hotspot V600E. *NF1* demonstrated a relatively non-selective mutation pattern across cancer types, with modest enrichment in SKCM. *NRAS*, in contrast, showed significant and specific enrichment in SKCM, with mutational hotspots at Q61R and Q61K. The previous TCGA study identified *BRAF, NRAS*, and *NF1* as key mutated genes in SKCM [34], and we extended these findings by showing that they are also specific to SKCM. *EGFR* exhibited selective mutation enrichment in GBM, followed by LUAD. Interestingly, *EGFR* mutations in GBM were concentrated at A289V and G598V, while *EGFR* mutations in LUAD were concentrated at L858R and E746_A750del. *ERBB2*, while well-known for amplification in breast cancer [44], showed only modest mutation frequencies across cancer types, with the highest occurrence in CAC. The findings revealed that genes participating in the RTK/RAS/MAPK pathway, including *EGFR, ERBB2, KRAS, NRAS, NF1*, and *BRAF* had highly heterogeneous mutation patterns. *APC* exhibited significant and specific mutations in CRC, a well-established defining feature of colorectal cancer [45]. *CTNNB1* showed frequent mutations in HCC and EC, followed by ACC. Despite both genes functioning within the WNT pathway, *APC* and *CTNNB1* demonstrated completely distinct mutation patterns. *RB1*, the well-known tumor suppressor gene, showed frequent mutations in BUC and LMS. *CDKN2A*, although more often altered by deletion, also exhibited frequent mutations in HNSCC, PDAC, and LUSC, with a mutational hotspot at R80*. These two cell cycle-related genes demonstrated distinct mutation patterns.

*KMT2D*, a gene involved in chromatin regulation, was widely mutated with the highest frequencies in BUC, EC, and LUSC, all of which are characterized by high background mutation burdens [8]. Given *KMT2D*’s extensive length (>5,500 amino acids [46]) and absence of recurrent mutational hotspots, these alterations were likely passenger mutations rather than functional drivers. *ARID1A*, another gene related to chromatin regulation, also showed widespread mutations with the highest frequencies in EC, followed by GAC and BUC. Additional genes involved in chromatin regulation include *KMT2C, EP300*, and *KDM6A. KMT2C* showed widespread mutations similar to *KMT2D*, with the highest frequencies in CAC, CSCC, and BUC. *EP300* exhibited frequent mutations in BUC and CSCC. *KDM6A* demonstrated highly specific mutations in BUC. Collectively, these findings revealed that chromatin regulation genes showed preferential mutations in BUC, consistent with previous TCGA findings [25], and also suggest that these mutations may not be purely passenger events. *FAT1* exhibited a widespread mutation pattern but was particularly enriched in HNSCC, followed by EC. *NOTCH1* showed frequent mutations in HNSCC, ESCC, and ODG. Both of these transmembrane proteins involved in signaling between adjacent cells were enriched in HNSCC. *ATM*, a tumor suppressor gene frequently associated with germline mutations especially in PDAC [47], showed widespread and non-selective somatic mutations. *BRCA2*, known for its association with breast and ovarian cancer through germline mutations [47], also exhibited widespread and non-selective somatic mutations. These findings suggest that although germline mutations of these DNA repair genes are cancer-type-specific, their somatic mutations are not. Several genes demonstrated frequent frameshift mutations, including *RNF43, MSH3, JAK1, BCORL1, CTCF*, and *MSH6*, with these mutations being most prevalent in EC, followed by GAC and CRC. This pattern revealed the existence of microsatellite instability (MSI) subtypes of cancers, characterized by defects in DNA mismatch repair that lead to exceptionally frequent frameshift mutations at short tandem repeats of DNA [48] [49]. The identification of MSI tumors in EC, GAC, and CRC was consistent with previous TCGA studies [30] [14] [5]. *KEAP1* exhibited frequent mutations in LUAD. *NFE2L2* showed frequent mutations in ESCC and LUSC. Both genes are involved in protection against oxidative stress and were enriched in cancers of the aerodigestive tracts. *SMAD4*, a mediator of the TGFβ pathway, was significantly mutated in PDAC.

*IDH1* mutations, predominantly at hotspot R132H and leading to accumulation of the oncometabolite 2HG [50], were highly significant and specific to lower-grade gliomas, reflecting a defining molecular feature of lower-grade gliomas associated with better prognosis [38]. *IDH1* was also mutated in CCA, though to a lesser extent. *ATRX* exhibited significant mutations in lower-grade gliomas, particularly in AC and OAC, consistent with the *IDH*-mutated without 1p/19q codeletion subtype identified in the TCGA study [38]. *ATRX* was also frequently mutated in soft tissue sarcomas, especially UPS. *CIC* was significantly mutated in ODG, followed by OAC, consistent with the *IDH*-mutated with 1p/19q codeletion subtype identified in the TCGA study [38]. *FUBP1* showed a similar but weaker pattern to *CIC*. Together, the mutational status of these genes forms the molecular foundation for lower-grade glioma subtype classification. Several genes were selectively mutated in SKCM, including *PTPRT, ROS1, DPYD, SNCAIP, ERBB4*, and *TP63*, which were not highlighted in the previous TCGA study [34]. While the diverse functions of these genes and the high mutational burden of SKCM might suggest these are passenger events, their significantly shorter amino acid lengths combined with higher mutation frequencies compared to *KMT2D* may indicate potential biological significance. *FBXW7* exhibited frequent mutations in UCS, followed by EC and CRC. *PPP2R1A* showed frequent mutations in UCS, followed by EC. Both genes demonstrated preferential enrichment in UCS. *VHL* exhibited significant and specific mutations in ccRCC, representing a defining molecular feature of this cancer type [22]. *PBRM1* showed frequent and selective mutations in ccRCC, with additional involvement in CCA. *SETD2* was moderately mutated in ccRCC. *BAP1* demonstrated frequent and selective mutations in MPM, UM, and CCA, representing a characteristic feature of MPM [37] and the monosomy 3 subtype of UM [35], while also being mildly mutated in ccRCC. Notably, all four genes frequently mutated in ccRCC are located on the short arm of chromosome 3 [22]. *CDH1* exhibited specific mutations in ILC, representing the hallmark of this cancer type that leads to decohesive neoplastic cells [33]. *GATA3* showed elevated mutations in IDC, associated with the luminal A subtype [4]. *SF3B1* demonstrated selective mutations in UM, representing a characteristic feature of the disomy 3 subtype [35]. *KIT* exhibited frequent mutations in TGCT, a molecular feature of seminioma [28]. *GTF2I* showed specific mutations in THYM at mutational hotspot L424H, representing the hallmark of this cancer type [42]. *NF2* demonstrated selective mutations in MPM, a characteristic feature of this malignancy [37]. *STK11* exhibited elevated mutations in LUAD, leading to mTOR pathway activation [12]. *GNAQ* and *GNA11* represented the defining molecular alterations of UM [35], with mutational hotspots at Q209L for both genes and Q209P for GNAQ. Several genes showed specific mutations in DLBCL, including *BTG2, B2M, CARD11, PIM1, KLHL6, P2RY8*, and *SOCS1*. Additionally, *KMT2D* was disproportionately mutated in DLBCL.

### Copy Number Variation Analysis

We analyzed the copy number variation data to characterize cancer-type-specific gene amplification and deletion patterns. Log-transformed chromosomal segment values were converted back to copy numbers, and gene copy numbers were determined based on their containing segments. The heatmap displays arm-level copy number changes across cancer types for each sample (Figure 3). The bubble plots show amplification (left) and deletion (right) frequencies for genes exhibiting focal copy number alterations or having established biological relevance (Figure 4).

**Figure 3:**
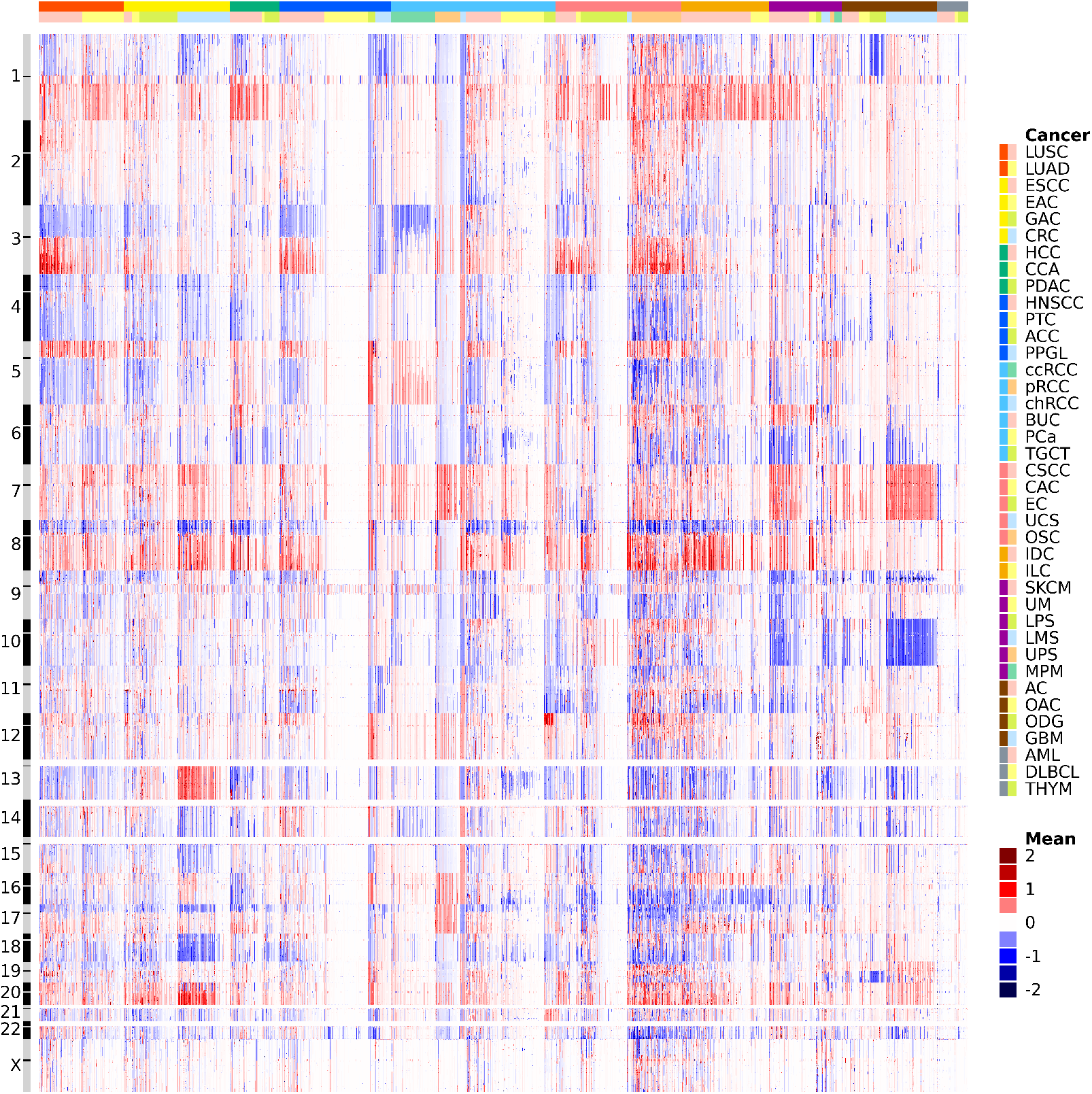
Copy number landscapes across cancer types. Chromosomal copy number variations across cancer samples. Color intensity represents the log-transformed fold change in copy number, with red indicating gains and blue indicating losses. Chromosome numbers and centromere positions are shown in the left sidebar; cancer types are indicated in the top color bar.

**Figure 4:**
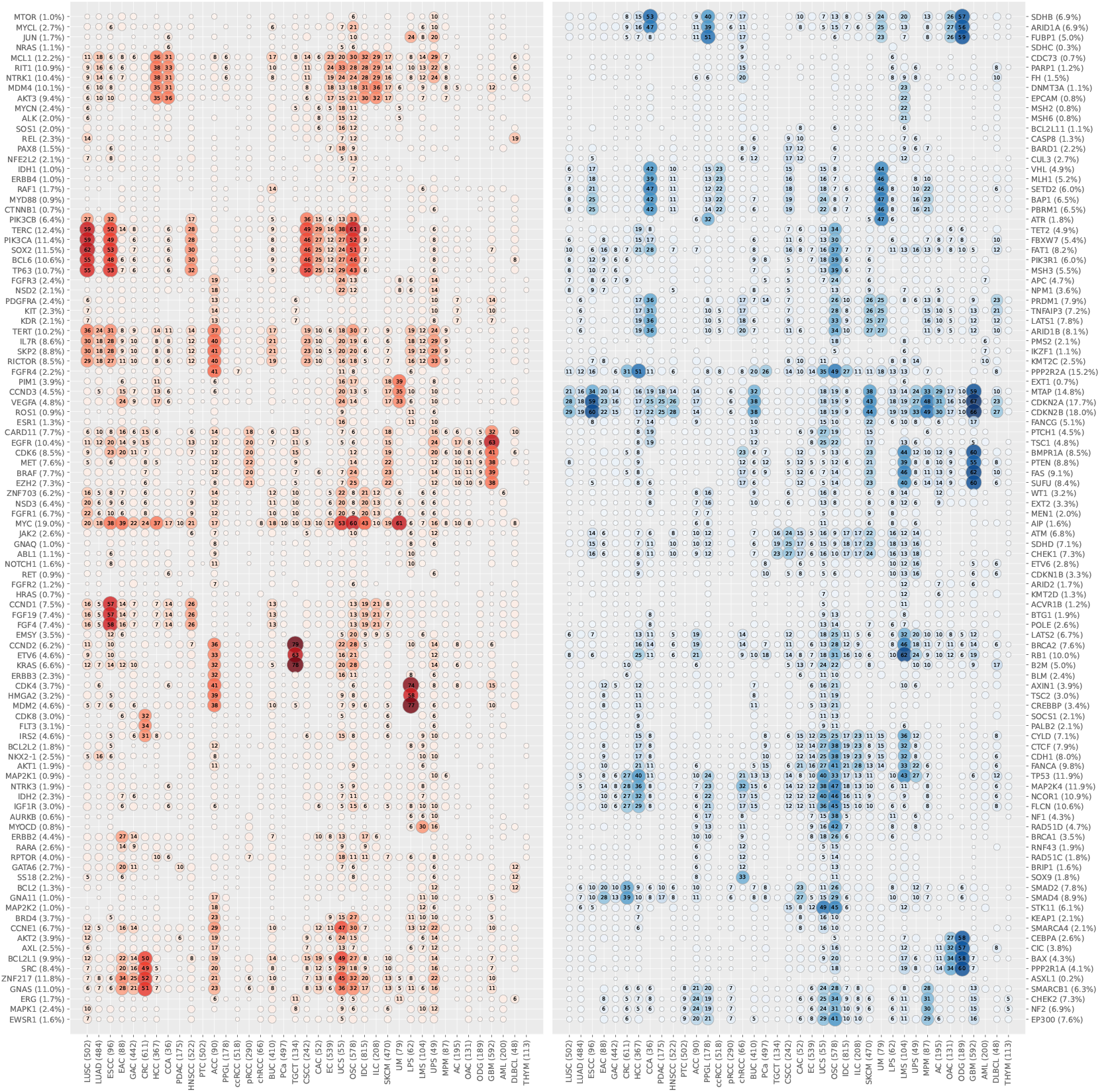
Gene amplifications and deletions across cancer types. Amplification frequencies (left) and deletion frequencies (right) for tumor-related genes across cancer types. Bubble size and color intensity represent alteration frequency. Genes are ordered by chromosomal position. Numbers following gene names indicate overall alteration frequencies; numbers following cancer type names indicate sample sizes.

The majority of cancer types demonstrated consistent arm-level amplification or deletion patterns, suggesting that the selective advantage of specific chromosomal arm alterations is shared across different cancer types. Several cancer types exhibited genomically stable profiles with minimal copy number changes, including PTC, PCa, a subset of EC, and AML. ChRCC displayed a distinctive pattern characterized by frequent whole-chromosomal alterations, with gains of chromosomes 4, 7, 11, 12, 14, 15, 16, 18, 19, 20, and 22, and losses of chromosomes 1, 2, 6, 10, 13, and 17 [51].

Chromosome 1p exhibited focal amplification of *JUN* in LPS and UPS, focal amplification of *MYCL* in OSC, arm-level deletions in ODG, OAC, PPGL, and UM, and terminal deletion in CCA. The 1p deletion in ODG is consistent with the characteristic 1p/19q codeletion pattern [38] and results in *FUBP1* deletion, which acts synergistically with *FUBP1* mutation characteristic of ODG. Chromosome 1q demonstrated arm-level amplification in HCC, CCA, IDC, ILC, and OSC, terminal amplification in UPS, focal amplification of *RIT1* in UCS, and focal deletion of *FH* in chRCC. Potential target genes of arm-level amplifications include *MCL1* and *MDM4*. Chromosome 2p exhibited focal amplification of *REL* in DLBCL and arm-level deletion in LMS. Chromosome 2q demonstrated focal amplification of *PAX8* in UCS. Chromosome 3p exhibited arm-level deletions in UM, CCA, ccRCC, and ESCC, and focal deletion of *BAP1* in MPM. Four frequently mutated genes in ccRCC—*VHL, SETD2, BAP1*, and *PBRM1* [22]—are all located on 3p, indicating synergistic effects between copy number loss and somatic mutations. Similarly, CCA harbors frequent *BAP1* and *PBRM1* mutations. Chromosome 3q demonstrated arm-level amplifications in LUSC, ESCC, HNSCC, CSCC, CAC, OSC, and UCS, and arm-level deletions in UM and PPGL. Potential target genes of arm-level amplifications include *SOX2* [52] and *TP63*, while *TERC* is the primary amplification target in OSC. The 3p and 3q deletions in UM are consistent with the monosomy 3 subtype associated with poor prognosis [35]. Chromosome 4p exhibited focal amplification of *FGFR3* in UCS. Chromosome 4q demonstrated terminal deletion in OSC. Chromosome 5p exhibited arm-level amplifications in LUSC, ESCC, CSCC, ACC, and UPS, as well as focal amplification of *TERT* in OSC. Potential target genes of arm-level amplifications include *TERT* and *RICTOR*. Chromosome 5q demonstrated arm-level amplification in ACC and focal deletion of *PIK3R1* in OSC.

Chromosome 6p exhibited arm-level amplifications in UM and focal amplifications of *VEGFA* in EAC and UCS. Chromosome 6q demonstrated arm-level deletions in CCA, OSC, SKCM, and UM, and focal deletions of *PRDM1* and *TNFAIP3* in DLBCL. Potential target genes of arm-level deletions include *LATS1* and *ARID1B*. Chromosome 7p exhibited arm-level amplification and focal amplification of *EGFR* in GBM. Chromosome 7q demonstrated arm-level amplifications in GBM, pRCC, and SKCM, terminal amplification in OSC, and focal amplifications of *CDK6* in ESCC and UPS. Potential target genes of arm-level amplifications include *CDK6* and *BRAF*. Chromosome 8p exhibited focal amplification of *FGFR1* in LUSC, focal amplifications of *ZNF703* in UCS and IDC, arm-level deletion in HCC, and terminal deletions in OSC, UCS, CRC, IDC, and PCa. Potential target genes of arm-level deletions include *PPP2R2A*. Chromosome 8q demonstrated arm-level amplifications in UM, UCS, OSC, IDC, and HCC, with focal amplifications of *MYC* in OSC, UCS, IDC, ESCC, and EAC. Chromosome 9p exhibited arm-level deletions in GBM and SKCM, and focal deletions of *CDKN2A* and *CDKN2B* in GBM, ESCC, MPM, SKCM, BUC, UPS, AC, LUSC, HNSCC, DLBCL, PDAC, CCA, and EAC. Chromosome 9q demonstrated arm-level deletion in UCS and OSC, with potential target genes including *TSC1*. Chromosome 10p and 10q exhibited arm-level deletions in GBM, LMS, chRCC, and SKCM, with potential target genes including *PTEN*.

Chromosome 11p exhibited terminal deletion in OSC. Chromosome 11q demonstrated focal amplifications of *CCND1* in ESCC, HNSCC, ILC, and IDC, focal amplification of *EMSY* in OSC, and terminal deletions in CSCC and SKCM. Potential target genes of terminal deletions include *ATM*. Chromosome 12p exhibited arm-level amplification in TGCT, ACC, OSC, and UCS. The characteristic 12p amplification in TGCT is related to isochromosome formation [53], with possible target genes including *KRAS*. Interestingly, *ETV6* showed relative deletion in TGCT compared to the highly amplified background. Chromosome 12q demonstrated arm-level amplification in ACC and focal amplifications of *MDM2* and *CDK4* in LPS, the defining feature of LPS [54]. Chromosome 13q exhibited arm-level amplification in CRC, arm-level deletions in LMS, UPS, and OSC, and focal deletion of *RB1* in LMS, OSC, and HCC. Potential target genes of arm-level amplifications include *CDK8* and *IRS2*, while the target gene of arm-level deletions is *RB1*. Chromosome 14q exhibited focal amplification of *NKX2-1* in LUAD and terminal deletion in CCA. Chromosome 15q exhibited terminal deletions in UCS and OSC, and focal deletion of *B2M* in DLBCL.

Chromosome 16p exhibited arm-level amplification in ACC and terminal deletion in OSC. Potential target genes include *CREBBP*. Chromosome 16q demonstrated arm-level amplification in ACC and arm-level deletions in OSC, LMS, UCS, and ILC. Potential target genes of arm-level deletions include *CDH1*. Chromosome 17p exhibited focal amplification of *MYOCD* in LMS and UPS, arm-level deletions in OSC, UCS, HCC, CRC, and PPGL, and focal deletion of *MAP2K4* in chRCC. *TP53* is the target gene of arm-level deletions in LMS, HCC, UCS, OSC, CRC, UPS, and PPGL. Chromosome 17q demonstrated focal amplifications of *ERBB2* in EAC and IDC, focal deletions of *NF1* and *RAD51D* in OSC, and focal deletion of *SOX9* in chRCC. The *ERBB2* amplifications in IDC and EAC provide the rationale for using HER2 inhibitors in these cancer types [55] [56]. Chromosome 18p exhibited arm-level deletion in CRC. Chromosome 18q demonstrated focal amplification of *GATA6* in EAC, arm-level deletion in CRC, and focal deletion of *SMAD4* in CRC, EAC, CAC, and OSC. Chromosome 19p exhibited arm-level amplification in ACC, focal amplification of *BRD4* in OSC, and terminal deletions in UCS and OSC. Chromosome 19q demonstrated arm-level amplification in ACC, focal amplification of *CCNE1* in UCS, OSC, and UPS, and arm-level deletions in ODG and OAC. Similar to 1p, the 19q deletion in ODG is consistent with the 1p/19q codeletion pattern and results in *CIC* deletion acting synergistically with *CIC* mutation in ODG. Chromosome 20p exhibited arm-level amplification in ACC. Chromosome 20q demonstrated arm-level amplifications in CRC, UCS, OSC, EAC, and ACC. Potential target genes of arm-level amplifications include *BCL2L1* and *ZNF217*. Chromosome 21q exhibited relatively stable copy number profiles across cancer types. Chromosome 22q exhibited arm-level deletions in OSC, UCS, MPM, and ACC. Potential target genes of arm-level deletions include *CHEK2* and *NF2*.

### DNA Methylation Analysis

We analyzed the DNA methylation data to characterize cancer-type-specific hypermethylation and hypomethylation patterns. Hypermethylated CpG sites were defined as those methylated in >30% of cancer samples while remaining unmethylated in >80% of normal samples. Hypomethylated CpG sites were defined vice versa. Hypermethylated genes were defined as having hypermethylated CpG sites in their promoter regions. Hypomethylated genes were defined vice versa. The bubble plots display the hypermethylation (left) and hypomethylation (right) frequencies of the top differentially methylated genes across cancer types (Figure 5).

**Figure 5:**
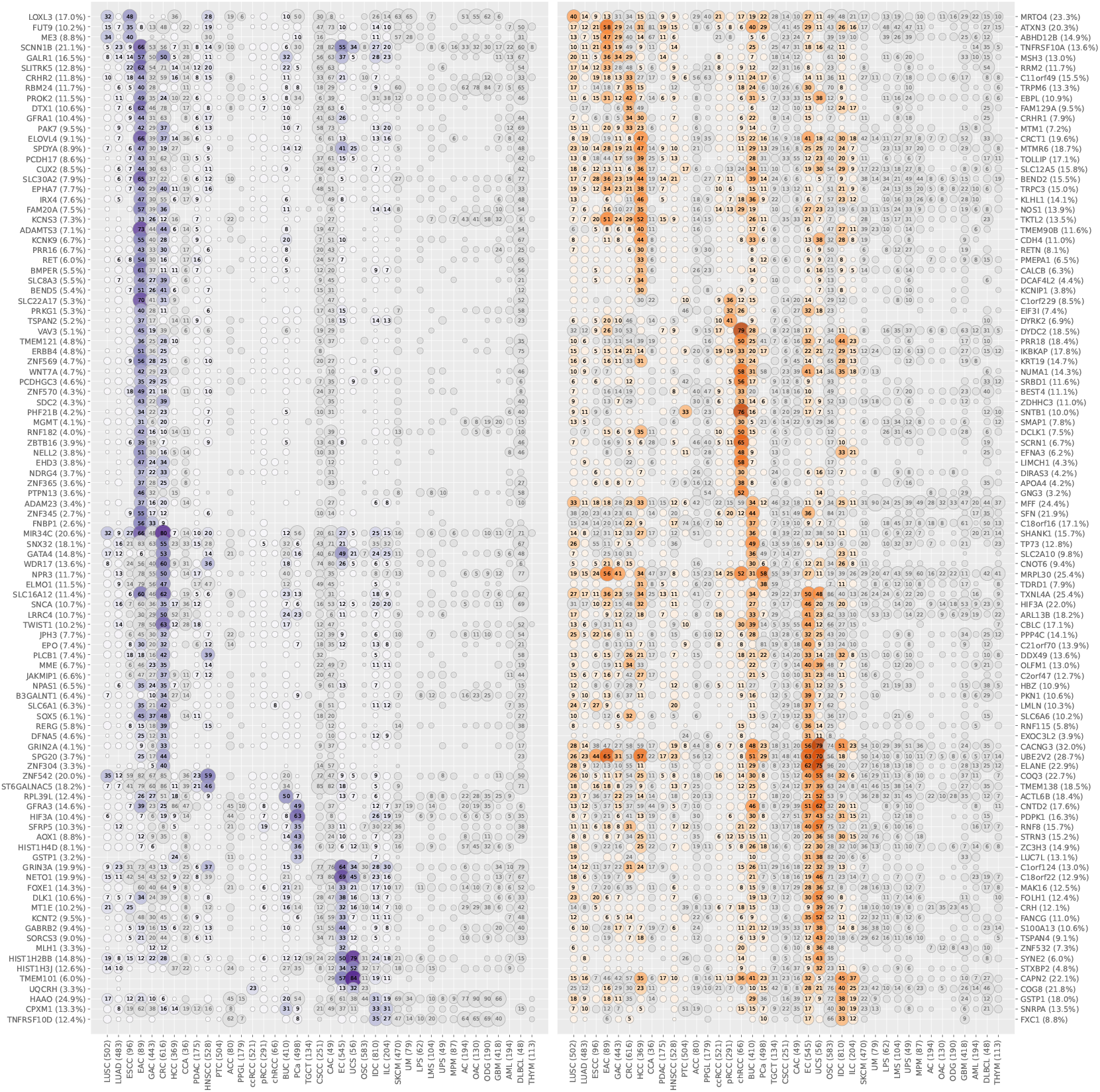
Gene hypermethylations and hypomethylations across cancer types. Hypermethylation frequencies (left) and hypomethylation frequencies (right) for differentially methylated genes across cancer types. Bubble size and color intensity represent alteration frequency. Gray bubbles indicate genes that do not meet the baseline methylation criteria in normal samples for specific cancer types. Genes are grouped by their most relevant cancer types and ordered by alteration frequency. Number following gene names indicate overall alteration frequencies; numbers following cancer type names indicate sample sizes.

Compared to simple nucleotide variation and copy number variation, DNA methylation exhibited more widespread and non-specific alteration patterns, with hypomethylation being more non-specific than hypermethylation. Nevertheless, our approach successfully identified three well-established cancer-associated hypermethylated genes: *GSTP1, MGMT*, and *MLH1. GSTP1* has long been recognized as a frequently hypermethylated gene in prostate cancer [57] and suggested to be implicated in early carcinogenesis [58]. Our analysis revealed that *GSTP1* hypermethylation is highly specific to prostate cancer, followed by hepatocellular carcinoma. *MGMT* encodes a DNA repair protein that removes alkyl adducts from guanine, protecting cancer cells from DNA damage caused by alkylating agents. Research has shown that *MGMT* is often hypermethylated in specific cancer types [59], and patients with glioblastoma containing methylated *MGMT* benefit from temozolomide [60]. Our results showed that besides GBM, *MGMT* was also significantly hypermethylated in EAC and CRC, suggesting that alkylating agent use should be investigated in these cancer types. *MLH1* is a core DNA mismatch repair gene whose hypermethylation leads to microsatellite instability in cancers, particularly colorectal [61] [62], endometrial [63], and gastric cancers [64]. Our analysis revealed that *MLH1* hypermethylation was most pronounced in EC, followed by GAC and CRC, which aligns with our SNV results showing that MSI subtypes are predominantly found in these three cancer types.

Additional hypermethylated genes with potential biological significance are summarized below. *PCDH17*, a protocadherin acting as a tumor suppressor inducing apoptosis and autophagy [65], was hypermethylated in EAC. *EPHA7*, a tumor suppressor for follicular lymphoma [66], was hypermethylated in EAC and CRC. *ZBTB16*, a transcription factor contributing to melanoma progression when silenced [67], was hypermethylated in EAC. *NDRG4*, a candidate tumor suppressor suppressing cell proliferation and invasion [68], was hypermethylated in EAC and CRC. *ADAM23*, whose silencing is correlated with tumor progression and metastasis [69], was hypermethylated in EAC. *MIR34B/C*, microRNAs that are transcriptional targets of *TP53* [70], were hypermethylated in CRC and EAC. *GATA4*, a transcription factor causing senescence in response to DNA damage [71], was hypermethylated in CRC and EC. *TWIST1*, a transcription factor promoting epithelial-mesenchymal transition (EMT) [72], was hypermethylated in CRC. *DFNA5*, a pore-forming protein that can switch apoptosis to pyroptosis under chemotherapy drugs [73], was hypermethylated in CRC. *SFRP5*, a Wnt signaling modulator whose epigenetic silencing contributes to malignancy and chemoresistance [74], was hypermethylated in PCa.

### Transcriptome Profiling Analysis

We analyzed the transcriptome profiling data to characterize cancer-type-specific gene set enrichment patterns. Single-sample gene set enrichment analysis (ssGSEA) [75] was performed on log-transformed mRNA expression values using MSigDB Hallmark gene sets [76], and average scores were calculated after normalization across gene sets and across all samples. The heatmap displays average enrichment scores for each gene set across cancer types (Figure 6).

**Figure 6:**
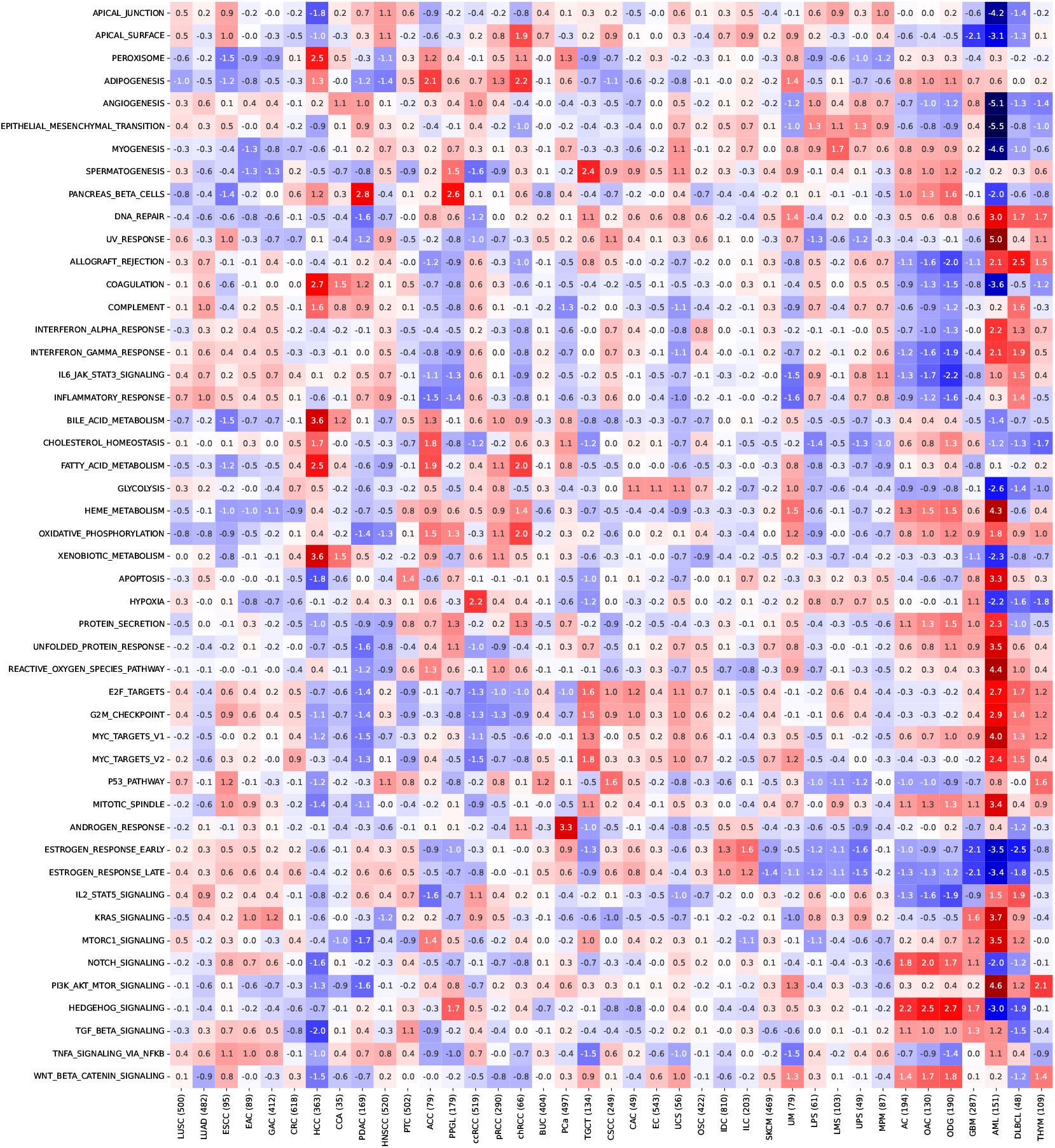
Hallmark gene set enrichment across cancer types. Single-sample gene set enrichment analysis (ssGSEA) scores for Hallmark gene sets across cancer types. Color intensity represents the Z-transformed average enrichment score, with red indicating positive enrichment and blue indicating negative enrichment. Numbers following cancer type names indicate sample sizes.

Most cancer types demonstrated significant enrichment in only a few gene sets, often reflecting their tissue of origin. AML presented as an outlier with disproportionately high enrichment values across most gene sets, warranting separate discussion. AML showed significant positive enrichment in KRAS signaling, PI3K/AKT/mTOR signaling, mTORC1 signaling, MYC targets, and mitotic spindle, suggesting a highly active and proliferative state; in apoptosis, unfolded protein response, reactive oxygen species pathway, DNA repair, and UV response, indicating the presence of constant cellular stress; and in heme metabolism, consistent with its hematopoietic origin. AML also showed significant negative enrichment in apical junction, apical surface, angiogenesis, epithelial-mesenchymal transition, and myogenesis, reflecting the absence of stromal components.

Apical surface genes demonstrated activation in chRCC, which may correspond to the microscopic finding of prominent eosinophilic cell membranes in chRCC [77]. Peroxisome genes showed significant activation in HCC, reflecting peroxisomal β-oxidation and detoxification in the liver. Adipogenesis genes exhibited significant activation in chRCC and ACC, but interestingly not in LPS. Epithelial-mesenchymal transition genes demonstrated activation in LPS and UPS, consistent with their mesenchymal origin. Myogenesis genes showed activation in LMS, consistent with its muscle origin. Spermatogenesis genes exhibited significant activation in TGCT, consistent with its germ cell origin, while also showing activation in PPGL. Pancreatic β-cell genes demonstrated significant activation in PDAC as expected, while also showing significant activation in PPGL, reflecting their shared neuroendocrine characteristics. DNA repair genes showed activation in DLBCL and THYM, in addition to AML. Allograft rejection genes exhibited significant activation in DLBCL, AML, and THYM, consistent with their immune system origin. Coagulation genes demonstrated significant activation in HCC and related CCA, reflecting the liver’s role in coagulation factor synthesis. Complement genes showed activation in HCC, reflecting the liver’s role in complement synthesis, while also showing activation in DLBCL. Interferon-α and interferon-γ response genes exhibited significant activation in AML and DLBCL, reflecting immune cells’ roles in producing and responding to cytokines. IL-6/JAK/STAT3 signaling and inflammatory response genes demonstrated activation in DLBCL, correlating with the systemic inflammatory response associated with lymphoma (B-symptoms) [78].

Bile acid metabolism genes demonstrated remarkable activation in HCC, consistent with the liver’s role in bile acid synthesis. Cholesterol homeostasis genes showed activation in ACC and HCC, consistent with the steroidogenic function of the adrenal cortex and the liver’s role in cholesterol metabolism. Fatty acid metabolism genes exhibited significant activation in HCC, chRCC, and ACC. Heme metabolism genes demonstrated activation in OAC, ODG, and UVM, in addition to AML. Oxidative phosphorylation genes showed significant activation in chRCC, in accordance with previous findings that chRCC demonstrates increased utilization of mitochondrial ATP generation [24], while activation was also observed in AML and ACC. Xenobiotic metabolism genes exhibited remarkable activation in HCC and related CCA, reflecting the liver’s central role in detoxification. Hypoxia genes demonstrated significant activation in ccRCC, reflecting activation of hypoxia-inducible factor pathways due to VHL inactivation in ccRCC [79]. Protein secretion genes showed significant activation in AML and ODG. E2F targets, G2/M checkpoint, and MYC targets genes exhibited coordinated and significant activation in AML, followed by DLBCL and TGCT. The elevated cell cycle pathways reflected the highly proliferative nature of hematologic malignancies and germ cell tumors, while the coordination with MYC targets suggested that MYC closely correlates with cell cycle progression [80]. Notably, c-MYC is a Yamanaka pluripotency factor [81], which is consistent with the pluripotent nature of germ cell tumors (TGCT). p53 pathway genes demonstrated activation in CSCC and THYM.

Androgen response genes demonstrated remarkable activation in PCa, consistent with the androgen-dependent nature of PCa [82]. Estrogen response genes showed activation in ILC and IDC, consistent with the hormone-dependent nature of many breast cancers. IL-2/STAT5 signaling genes exhibited activation in DLBCL and AML, reflecting the critical role of IL-2 signaling in the immune system. KRAS signaling genes demonstrated activation in GBM in addition to AML, while cancer types with frequent KRAS mutations, including PDAC and CRC, did not show activation. Notch signaling genes showed activation in AC, OAC, and ODG, reflecting the crucial role of Notch signaling in neural development [83]. PI3K/AKT/mTOR signaling genes exhibited significant activation in THYM in addition to AML. Hedgehog signaling genes demonstrated significant activation in ODG, OAC, AC, and GBM, suggesting a role for Hedgehog signaling in neural development similar to Notch signaling. However, in contrast to the inactivation of Notch signaling in PPGL, Hedgehog signaling showed activation in PPGL. Wnt/β-catenin signaling genes showed activation in ODG, OAC, AC, and THYM, but did not show activation in CRC, which harbors predominantly APC mutations.

### Multiomics Clustering Analysis

We performed unsupervised hierarchical clustering of mRNA expression, miRNA expression, protein expression, and DNA methylation data to characterize molecular relationships among cancer types. Log-transformed values of mRNA and miRNA expression, abundance values of protein expression, and beta values of DNA methylation were normalized before hierarchical clustering was performed across both samples and genes or CpG sites. The heatmaps display the clustering results from each molecular platform (Figure 7).

**Figure 7:**
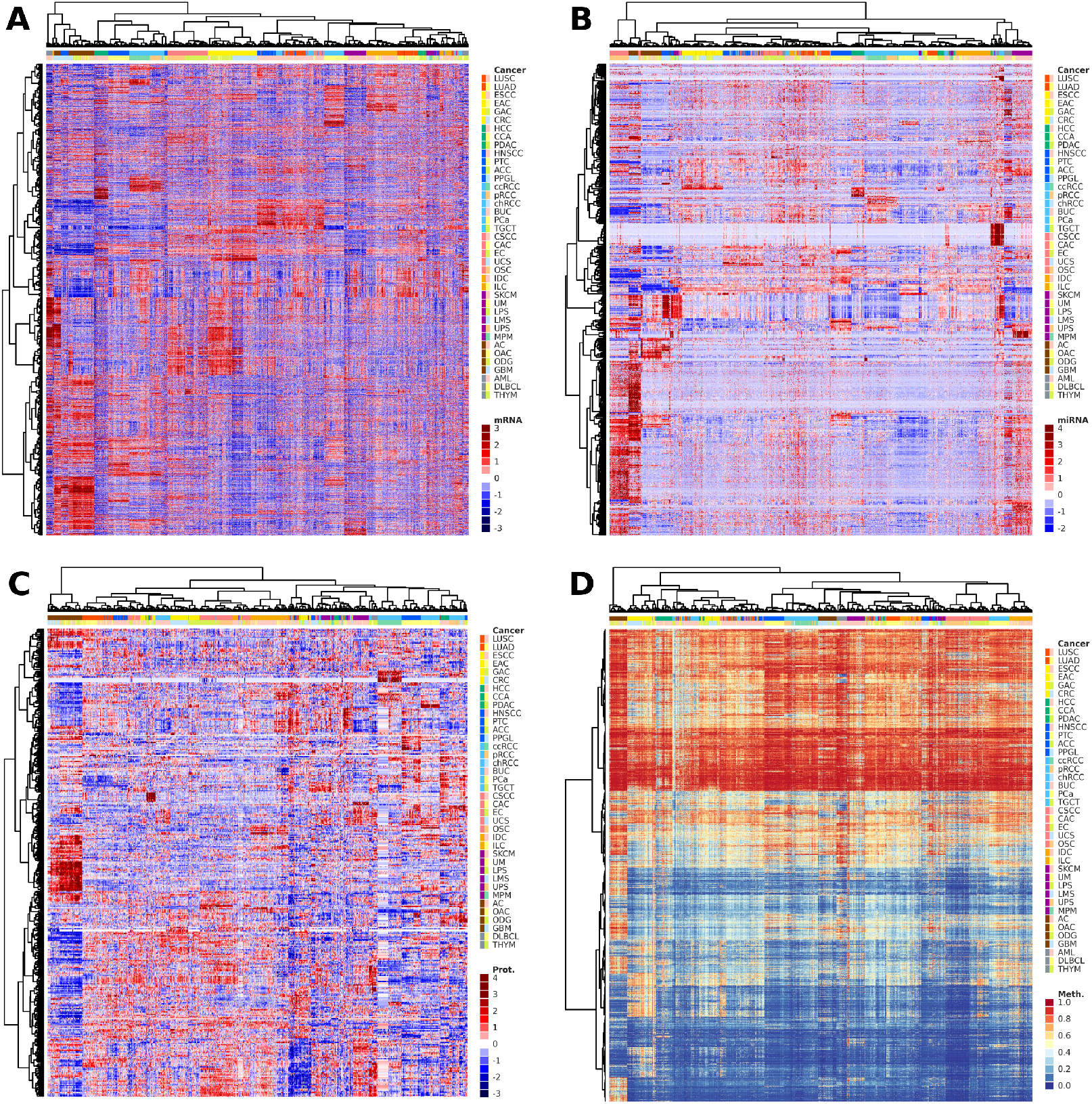
Unsupervised clustering of samples across cancer types. Unsupervised hierarchical clustering of (A) mRNA expression, (B) miRNA expression, (C) protein expression, and (D) DNA methylation data. Color intensity represents normalized expression or methylation values. The top color bar indicates cancer type, the top dendrogram displays sample clustering, and the left dendrogram displays gene or CpG site clustering.

mRNA expression clustering demonstrated the most consistent and biologically coherent results. Squamous cell carcinomas formed a distinct pan-squamous cluster comprising LUSC as a distinct cluster, HNSCC as a distinct cluster, BUC as two distinct clusters, and CSCC and CAC as two adjacent clusters. Gastrointestinal malignancies formed a distinct pan-gastrointestinal cluster containing ESCC as a distinct cluster, EAC and GAC as an intermixed cluster, and CRC as a distinct cluster. Renal and adrenal tumors formed a distinct pan-kidney cluster comprising ccRCC as a distinct cluster, pRCC as a distinct cluster, and chRCC and ACC as two adjacent clusters. Gynecological malignancies formed a distinct pan-gynecological cluster containing EC as a distinct cluster and OSC as a distinct cluster. Neural tumors formed a distinct pan-neural cluster containing AC, OAC, and ODG as an intermixed cluster, GBM as two distinct clusters, and PPGL as a distinct cluster. Sarcomas formed a distinct pan-sarcoma cluster comprising LPS and UPS as an intermixed cluster, LMS as a distinct cluster, and MPM as a distinct cluster.

Individual cancer types formed separate clusters including LUAD as a distinct cluster, HCC as a distinct cluster, CCA and PDAC as two adjacent clusters, PTC as a distinct cluster, PCa as a distinct cluster, UCS as a distinct cluster, TGCT as a distinct cluster, IDC and ILC as an intermixed cluster, SKCM and UM as two adjacent clusters, AML as a distinct cluster, and DLBCL and THYM as two adjacent clusters.

DNA methylation clustering largely recapitulated mRNA expression clustering with several differences. The pan-squamous cluster was intermixed rather than distinct. ESCC was incorporated into the pan-squamous cluster instead of the pan-gastrointestinal cluster. CCA clustered with HCC rather than PDAC. ACC was separated from the pan-kidney cluster. PPGL was separated from the pan-neural cluster. A subset of BUC samples was separated from the pan-squamous cluster. A subset of UCS samples was incorporated into the pan-gynecological cluster. UM was separated from SKCM. GBM was separated from the pan-neural cluster. THYM was clustered with AML rather than DLBCL. miRNA expression clustering and protein expression clustering also demonstrated consistent results, with samples from the same cancer type predominantly clustering together. However, pan-cancer clusters were less apparent compared to mRNA expression clustering and DNA methylation clustering.

## Discussion

Our simple nucleotide variation analysis revealed that frequently mutated genes can be either widespread across cancer types or cancer-type-specific. Cancer-type-specific genes may co-occur with other driver mutations or represent the sole driver gene in certain cancer types. Widespread mutated genes include *TP53* showing predilection toward squamous cell carcinomas and OSC, *PIK3CA* showing predilection toward gynecologic malignancies, and *KRAS* showing predilection toward adenocarcinomas. Cancer-type-specific mutated genes co-occurring with other drivers include *PTEN* in EC, *BRAF* and *NRAS* in SKCM, *EGFR* in GBM, *APC* in CRC, *CTNNB1* in HCC, *IDH1*/*ATRX*/*CIC* in gliomas (with *ATRX* enriched in AC and *CIC* enriched in ODG), *VHL* in ccRCC, and *CDH1* in ILC. Cancer-type-specific mutated genes representing sole drivers include *BRAF* in PTC, *GTF2I* in THYM, and *GNAQ*/*GNA11* in UM. Some functionally related genes demonstrated similar mutation patterns, including MSI-related genes (*RNF43, MSH3*) in EC/GAC/CRC, chromatin regulation genes (*KMT2D, KDM6A*) in BUC, and adjacent cell signaling genes (*FAT1, NOTCH1*) in HNSCC. Integrative pathway analysis revealed that while different cancer types may share common altered pathways, the specific genes affected often differ. Within the PI3K/AKT pathway, *PIK3CA* showed widespread mutations, while *PTEN* and *PIK3R1* mutations were specific to EC. Within the RTK/RAS/MAPK pathway, *EGFR* was specific to GBM, *ERBB2* was widespread, *KRAS* was widespread, *NRAS* was specific to SKCM, *NF1* was widespread, and *BRAF* was specific to PTC and SKCM. Within the WNT pathway, *APC* was specific to CRC, and *CTNNB1* was specific to HCC.

Our copy number variation analysis revealed that arm-level alterations can be either widespread across cancer types or cancer-type-specific, with potential target genes of coherent biological significance often identifiable [84]. Widespread arm-level alterations include chromosome 1q amplification (*MCL1*), 3p deletion (*VHL*), 5p amplification (*TERT*), 7p amplification (*EGFR*), 7q amplification (*BRAF*), 8p deletion, 8q amplification (*MYC*), 9p deletion (*CDKN2A*), 17p deletion (*TP53*), and 20q amplification (*BCL2L1*). Cancer-type-specific arm-level alterations include chromosome 1p deletion in ODG (*FUBP1*), 3q amplification in squamous cell carcinomas (*SOX2*), 7p/q amplification in GBM (*EGFR*), 10p/q deletion in GBM (*PTEN*), 12p amplification in TGCT, 13q amplification in CRC, 18p/q deletion in CRC, and 19q deletion in ODG (*CIC*). Some arm-level alterations involved whole-chromosome changes, most characteristically observed in chRCC. Focal copy number alterations also exhibited both widespread and cancer-type-specific patterns. Widespread focal alterations include *MYC* amplification and *CDKN2A* deletion. Cancer-type-specific focal alterations include *JUN* amplification in LPS, *TERT* amplification in OSC, *VEGFA* amplification in EAC, *EGFR* amplification in GBM, *CCND1* amplification in ESCC, *MDM2* and *CDK4* amplification in LPS, *RB1* deletion in LMS, *MYOCD* amplification in LMS, *ERBB2* amplification in EAC, and *SMAD4* deletion in CRC. Integration of SNV and CNV analyses demonstrated that gene deletions frequently act synergistically with gene mutations, including *FUBP1* (1p) and *CIC* (19q) in ODG, *VHL* (3p) and adjacent genes in ccRCC, and *TP53* (17p) in various cancer types.

Our DNA methylation analysis identified several well-established cancer-associated hypermethylated genes, including *GSTP1* in PCa and HCC, *MGMT* in GBM and EAC/CRC, and *MLH1* in EC/GAC/CRC, consistent with our SNV results. Our analysis also revealed additional hypermethylated genes with potential biological significance, including *PCDH17* in EAC, *EPHA7* in EAC/CRC, *ZBTB16* in EAC, *NDRG4* in EAC/CRC, *ADAM23* in EAC, *MIR34B/C* in CRC/EAC, *GATA4* in CRC/EC, *TWIST1* in CRC, *DFNA5* in CRC, and *SFRP5* in PCa.

Our transcriptome profiling analysis revealed that cancer-type-specific pathway enrichment in cancer cells often reflects the normal physiological functions of their tissue of origin. For example, genes related to bile acid metabolism, fatty acid metabolism, cholesterol homeostasis, xenobiotic metabolism, peroxisome, coagulation, and complement were activated in HCC; androgen response genes in PCa; estrogen response genes in IDC and ILC; spermatogenesis genes in TGCT; myogenesis genes in LMS; pancreatic β-cell genes in PDAC; hypoxia genes in ccRCC; and IL-2 signaling, interferon response, and allograft rejection genes in AML and DLBCL. Other pathways showed significant enrichment in specific cancer types with less intuitive connections to tissue of origin. Genes related to apical surface and oxidative phosphorylation were activated in chRCC; cholesterol homeostasis genes in ACC; adipogenesis and fatty acid metabolism genes in chRCC and ACC; MYC target genes in TGCT; pancreatic β-cell genes in PPGL; and Notch signaling, Hedgehog signaling, and WNT signaling genes in gliomas (AC/OAC/ODG). Additionally, AML exhibited significant enrichment of genes associated with active proliferative states and persistent cellular stress.

Our multiomics clustering analysis identified several multi-cancer clusters composed of individual clusters from related cancers, most prominently in mRNA expression clustering. The pan-squamous cluster contained LUSC, HNSCC, BUC, CSCC, and CAC clusters. The pan-gastrointestinal cluster contained ESCC, EAC/GAC, and CRC clusters. The pan-kidney cluster contained ccRCC, pRCC, chRCC, and ACC clusters. The pan-gynecological cluster contained EC and OSC clusters. The pan-neural cluster contained AC/OAC/ODG, GBM, and PPGL clusters. The pan-sarcoma cluster contained LPS/UPS, LMS, and MPM clusters. Individual cancer type clusters included LUAD, HCC, CCA, PDAC, PTC, PCa, UCS, TGCT, IDC/ILC, SKCM, UM, AML, DLBCL, and THYM. DNA methylation clustering largely recapitulated mRNA expression clustering, while miRNA expression and protein expression clustering exhibited less distinct multi-cancer groupings.

In conclusion, we have provided a comprehensive atlas of cancer-type-specific molecular features through comparative analysis of TCGA data. The main contribution is our novel approach to identifying cancer-type-specific molecular features through unified comparative analysis and clear visualization that serves as a reference for hypothesis generation and clinical application. The main limitation is the primarily descriptive and correlative nature of this analysis, which lacks experimental validation. Future directions include experimental investigation of the underlying mechanisms and translation of these insights into clinical practice.

## Methods

### Data Acquisition

We obtained molecular data from The Cancer Genome Atlas (TCGA) via the Genomic Data Commons (GDC) API endpoints. File lists were queried from the file endpoint (https://api.gdc.cancer.gov/files/) by filtering on project IDs and data types, and data files were downloaded from the data endpoint (https://api.gdc.cancer.gov/data/). Samples were classified as tumor, normal (solid tissue normal), or blood (blood derived normal) based on their sample type annotations. Clinical data for all cases were downloaded directly from the GDC data portal (https://portal.gdc.cancer.gov). Additionally, we obtained leukocyte methylation data from the Gene Expression Omnibus (GSE35069) [10]. We analyzed 33 TCGA cancer projects: TCGA-ACC, TCGA-BLCA, TCGA-BRCA, TCGA-CESC, TCGA-CHOL, TCGA-COAD, TCGA-DLBC, TCGA-ESCA, TCGA-GBM, TCGA-HNSC, TCGA-KICH, TCGA-KIRC, TCGA-KIRP, TCGA-LAML, TCGA-LGG, TCGA-LIHC, TCGA-LUAD, TCGA-LUSC, TCGA-MESO, TCGA-OV, TCGA-PAAD, TCGA-PCPG, TCGA-PRAD, TCGA-READ, TCGA-SARC, TCGA-SKCM, TCGA-STAD, TCGA-TGCT, TCGA-THCA, TCGA-THYM, TCGA-UCEC, TCGA-UCS, and TCGA-UVM. Data types included masked somatic mutation (simple nucleotide variation), copy number segment (copy number variation), methylation beta value (DNA methylation), gene expression quantification (mRNA expression), miRNA expression quantification (miRNA expression), and protein expression quantification (protein expression).

### Cancer Cohort

We selected cases for each cancer type based on primary diagnoses recorded in the clinical data. Inclusion criteria followed standard pathological classification practices [77]. Our cohort comprised 39 cancer types: lung squamous cell carcinoma (LUSC), lung adenocarcinoma (LUAD), esophageal squamous cell carcinoma (ESCC), esophageal adenocarcinoma (EAC), gastric adenocarcinoma (GAC), colorectal adenocarcinoma (CRC), hepatocellular carcinoma (HCC), cholangiocarcinoma (CCA), pancreatic ductal adenocarcinoma (PDAC), head and neck squamous cell carcinoma (HNSCC), thyroid papillary carcinoma (PTC), adrenocortical carcinoma (ACC), pheochromocytoma and paraganglioma (PPGL), clear cell renal cell carcinoma (ccRCC), papillary renal cell carcinoma (pRCC), chromophobe renal cell carcinoma (chRCC), bladder urothelial carcinoma (BUC), prostate adenocarcinoma (PCa), testicular germ cell tumor (TGCT), cervical squamous cell carcinoma (CSCC), cervical adenocarcinoma (CAC), endometrial carcinoma (EC), uterine carcinosarcoma (UCS), ovarian serous carcinoma (OSC), breast invasive ductal carcinoma (IDC), breast invasive lobular carcinoma (ILC), cutaneous melanoma (SKCM), uveal melanoma (UM), liposarcoma (LPS), leiomyosarcoma (LMS), undifferentiated pleomorphic sarcoma (UPS), pleural mesothelioma (MPM), astrocytoma (AC), oligoastrocytoma (OAC), oligodendroglioma (ODG), glioblastoma (GBM), acute myeloid leukemia (AML), diffuse large B-cell lymphoma (DLBCL), and thymoma (THYM).

### Simple Nucleotide Variation Analysis

We defined tumor-related genes as a set containing 460 genes curated by the Japanese Cancer Genome Atlas (JCGA) [43] and 6 additional genes curated by the authors, including FOXP1, PAX8, LATS1, LATS2, GTF2I, and MYOCD. We extracted mutation annotations from each SNV file and excluded files with zero or more than 10,000 mutations. We retained only nonsynonymous mutations and categorized them into missense mutations, in-frame indels, critical site mutations (including splice site, splice region, translation start site, and nonstop mutations), frameshift indels, and nonsense mutations. We created waterfall plots for each cancer type, where each row represented a gene and each column represented a sample. In each waterfall plot, the top 20 tumor-related genes with the highest mutation frequencies were included, samples were ordered using lexicographical sorting of binary mutation vectors, and mutations were color-coded by category. We created bubble plots to compare mutation frequencies of genes and hotspots among different cancer types, where each row represented a gene or hotspot and each column represented a cancer type. We calculated mutation frequencies of genes across all samples and within each cancer type, and included tumor-related genes with mutation frequencies >10% in any cancer type, as well as the top remaining genes with the highest overall frequencies. In the bubble plot of genes, the 100 genes were ordered by overall frequency, and mutation frequencies were represented by the labels, sizes, and colors of bubbles. We also calculated mutation frequencies of hotspots across all samples and within each cancer type, and included hotspots of tumor-related genes that accounted for >5% of mutations in the gene, as well as the remaining top hotspots with the highest overall frequencies. In the bubble plot of hotspots, the 100 hotspots were ordered by overall frequency, while hotspots from the same gene were grouped together and frameshift mutations were moved to the end.

### Copy Number Variation Analysis

We extracted copy number segments from each CNV file and calculated CNV burdens, defined as the length-weighted averages of the absolute values of segment means. We created a heatmap to compare arm-level amplifications and deletions among different cancer types, where each row represented a chromosomal position, from chromosome 1 to chromosome X, and each column represented a sample, grouped by cancer type and ordered by CNV burden. All segments of each sample were mapped onto the heatmap based on their chromosomal positions and color-coded by their segment means, where red indicated amplifications and blue indicated deletions. Each cancer type was labeled with a unique color pair. We converted segment-level copy numbers to gene-level copy numbers to analyze gene-level amplifications and deletions. We used a gene-level copy number file from TCGA as the reference for start and end positions of all genes and implemented a two-pointer algorithm to assign each gene a copy number, defined as 2^1+*s*^ where *s* represents the segment mean of the segment containing the gene. We defined gene amplifications as copy number ≥ 2.83 (2^1.5^) and gene deletions as copy number ≤1.41 (2^0.5^), while excluding genes on chromosomes X and Y. We created bubble plots to compare gene-level amplifications and deletions among different cancer types, where each row represented a gene and each column represented a cancer type. We calculated amplification and deletion frequencies across all samples and within each cancer type, and included tumor-related genes that were substantially altered in any cancer type and had biological significance based on current knowledge. In the bubble plots of amplifications and deletions, the 100 genes were ordered by chromosomal position, and alteration frequencies were represented by the labels, sizes, and colors of bubbles.

### DNA Methylation Analysis

We defined target CpG sites as probes that were measured in all TCGA methylation files and located within promoter regions. We used three methylation beta value files from TCGA obtained via HM27, HM450, and EPIC arrays, and identified the intersection of the three sets of probes. We downloaded the annotation file for the HM450 array from the Illumina official website (https://sapac.support.illumina.com) and retained probes in the intersection that were labeled as TSS200 or TSS1500 for transcription start sites and labeled as island, N-shore, or S-shore for CpG island relationships. We extracted beta values from each methylation file, including tumor, normal, and leukocyte samples. We identified hypermethylation sites based on the following criteria. First, for each cancer type, we selected CpG sites that were unmethylated (defined as beta values ≤0.1) in more than 80% of normal samples from that organ but methylated (defined as beta values ≥0.5) in more than 30% of tumor samples from that cancer. Second, we retained only CpG sites that were unmethylated in more than 80% of overall normal samples and unmethylated in more than 80% of leukocyte samples. We identified hypomethylation sites similarly with individual and overall criteria, selecting CpG sites that were methylated (defined as beta values ≥ 0.5) in more than 80% of normal samples but unmethylated (defined as beta values ≤0.3) in more than 30% of tumor samples. Hypermethylation and hypomethylation sites were converted to hypermethylation and hypomethylation genes by mapping individual sites to their associated genes and retaining genes labeled as TSS200 or TSS1500. Only the site with the highest methylation or unmethylation percentage was retained if a gene corresponded to multiple sites. We created bubble plots to compare gene hypermethylation and hypomethylation among different cancer types, where each row represented a gene and each column represented a cancer type. We calculated the methylation and unmethylation frequencies of tumor and normal samples across all samples and within each cancer type, and included genes with frequencies >50% in any cancer type, as well as the top remaining genes with the highest overall frequencies. In the bubble plots of hypermethylation and hypomethylation, the 100 genes were grouped by the cancer types with the highest frequencies and ordered by methylation or unmethylation frequency. Methylation and unmethylation frequencies were represented by the labels, sizes, and colors of bubbles, where purple indicated hypermethylation and orange indicated hypomethylation, while grey indicated not being unmethylated or methylated in more than 80% of normal samples from that cancer type.

### Transcriptome Profiling Analysis

We downloaded the Hallmark gene set from MSigDB (https://www.gsea-msigdb.org/gsea/msigdb) to perform enrichment analysis [76]. We extracted TPM values from each RNA file and applied log_2_(*x* + 1) transformation. We performed single-sample gene set enrichment analyses (ssGSEA) following the original paper [75]. For each sample and each gene set, we sorted and normalized the expression values and calculated the running sum in descending order, which increased at target genes by values weighted by *x*^0.75^ for a total increase of 1 and decreased at non-target genes by equally weighted values for a total decrease of 1. The enrichment scores were defined as the integral of the running sum, calculated by summing each decreasing segment using the trapezoidal rule. For gene sets with both upregulated and downregulated subsets, we combined them by subtracting the downregulated scores from the upregulated scores. We calculated the enrichment scores for each cancer type on each gene set by first normalizing each sample across all gene sets, then normalizing each gene set across all samples, and finally calculating the average scores for each cancer type on each gene set. We created a heatmap to compare enrichment scores among different cancer types, where each row represented a gene set and each column represented a cancer type, and enrichment scores were represented by labels and colors.

### Multiomics Clustering Analysis

We performed hierarchical clustering on mRNA expression, miRNA expression, protein expression, and DNA methylation data. For mRNA expression data, we extracted raw expression counts from each mRNA file and retained genes that were expressed in more than half of the samples. We calculated geometric means for each gene across all samples while excluding zeros. We normalized each sample using size factors, which were the medians of the scaling factors for all genes, defined as the ratios of expression values to geometric means. We applied log_2_(*x* + 1) transformation and retained the top 20,000 genes with the highest variances. For miRNA expression data, we extracted RPM values from each miRNA file, applied log_2_(*x* + 1) transformation, and retained the top 1,000 genes with the highest variances. For protein expression data, we extracted relative abundance values from each protein file and retained the top 400 genes with the highest variances. For DNA methylation data, we extracted beta values from each file and excluded CpG sites on chromosomes X and Y and CpG sites with more than 10% missing values. We retained the top 10,000 CpG sites with the highest variances. Subsequent processes were identical for all data types. With the exception of DNA methylation data, we calculated z-scores for each gene across all samples. We performed hierarchical clustering on all cases and on all genes or sites, both using the standardized Euclidean distance and Ward’s method [85]. We created heatmaps with dendrograms to visualize the clustering results, where the scores of genes or sites were represented by colors, and cancer types were represented by color pairs.

### Code Implementation

The entire analysis workflow was implemented de novo by the authors, primarily in Python. Various Python packages were used, including Pandas for data manipulation [86], NumPy and SciPy for numerical computations [87] [88], and Matplotlib and Seaborn for data visualization [89] [90].

## Data Availability

All raw data files are publicly available through the GDC data endpoint (https://api.gdc.cancer.gov/data/). The processed metadata and summary files generated in this study are publicly available at https://github.com/hikarimusic/TCGA-Analysis.

## Acknowledgments

We thank the TCGA Research Network for making their data freely available. We also thank Reinius et al. for providing DNA methylation data of normal leukocytes, and the JCGA research team for curating tumor-related genes.

